# Cross-Sectional and Longitudinal Brain Scans Reveal Accelerated Brain Aging in Multiple Sclerosis

**DOI:** 10.1101/440412

**Authors:** Einar A. Høgestøl, Tobias Kaufmann, Gro O. Nygaard, Mona K. Beyer, Piotr Sowa, Jan E. Nordvik, Knut Kolskår, Geneviève Richard, Ole A. Andreassen, Hanne F. Harbo, Lars T. Westlye

**Author notes:** Corresponding author: Einar A. Høgestøl Department of Neurology, Neuroscience Research Unit, Multiple Sclerosis Research Group, University of Oslo / Oslo University Hospital, Domus Medica 4, room L-268, Gaustadalleén 34 0372 Oslo, Norway.

## Abstract

Multiple sclerosis (MS) is an inflammatory disorder of the central nervous system. By combining longitudinal MRI-based brain morphometry and brain age estimation using machine learning, we tested the hypothesis that MS patients have higher brain age relative to chronological age than healthy controls (HC) and that longitudinal rate of brain aging in MS patients is associated with clinical course.

Seventy-six MS patients, 71 % females and mean age 34.8 years (range 21-49) at inclusion, were examined with brain MRI at three time points with a mean total follow up period of 4.4 years. A machine learning model was applied on an independent training set of 3208 HC, estimating individual brain age and calculating the difference between estimated brain age and chronological age, termed brain age gap (BAG). We also assessed the longitudinal change rate in BAG in MS individuals. We used additional cross-sectional MRI data from 235 HC for case-control comparison.

MS patients showed increased BAG (4.4 ±6.6 years) compared to HC (Cohen’s D = 0.69, p = 4.0 × 10^−6^). Longitudinal estimates of BAG in MS patients suggested an accelerated rate of brain aging corresponding to an annual increase of 0.41 (±1.23) years compared to chronological aging for the MS patients (p = 0.008).

On average, patients with MS have significantly higher BAG compared to HC and accelerated rate of brain aging compared to chronological aging. Brain age estimation represents a promising method for evaluation of brain changes in MS, with potential for predicting future outcome and guide treatment.

## Abbreviations

BAG = Brain Age Gap

CNS = Central Nervous System

DMT = Disease-Modifying Treatment

EDSS = Expanded Disability Status Scale

FLAIR = Fluid Attenuation Inversion Recovery

FOV = Field of View

FSPGR = Fast Spoiled Gradient Echo

HC = Healthy Controls

ICC = Intraclass Correlation Coefficient

IQ = Intelligence Quotient

LME = Linear Mixed Effects

MP-RAGE = Magnetization Prepared Rapid Gradient Echo

MS = Multiple Sclerosis

MSSS = Multiple Sclerosis Severity Scale

NEDA = No Evidence of Disease Activity

TE = Echo Time

TR = Repetition Time

WMLL = White Matter Lesion Load

## INTRODUCTION

Multiple sclerosis (MS) is an inflammatory, demyelinating disease of the central nervous system (CNS). The prevalence of MS varies across the world and has been reported above 200/100.000 in some European countries ^1^. The disease affects women more frequently than men ^2^. The complex pathophysiology of MS can be divided into acute inflammation during a relapse and chronic inflammation thought to continuously disturb neuroaxonal homeostasis and drive neurodegeneration ^3^.

After the initial diagnosis, which is usually established in early adulthood, patients experience different disease activity and longitudinal accumulation of neurological damage. Development of robust brain imaging markers that can parse this between-subject heterogeneity of the clinical trajectories, predict future progression of disability, and monitor the effects of treatment is a major aim with important clinical implications ^4 5^. Current imaging markers with relevance for MS are associated with disease activity and progression, and include, among other features, number or volume of hyperintense brain lesions visible on T2-weighted MRI images, contrast-enhancing T1 lesions, increased annual brain volume loss and T1-hypointense “black holes” ^4 6 7^ Increased rate of brain volume loss, which is best captured using longitudinal designs ^8^, reflects accelerated neurodegeneration ^9^. Tuning the analysis towards specific brain regions may boost the correlations between estimated brain atrophy and disability ^4^. However, identifying robust associations between clinical outcomes and MRI measures has been challenging ^10^. This clinico-radiological paradox in MS is likely explained by a combination of lack of sensitivity and specificity both in the clinical and imaging domain. Indeed, expanded disability status scale (EDSS) and relapse rate, which are frequently used as clinical measures in MS phase III trials, are not sensitive to the full clinical spectrum in MS ^11 12^. Ongoing efforts are made to develop and validate novel multidimensional measures to capture subtle changes in MS activity and progression ^12^. The concept of “No evidence of disease activity” (NEDA) has emerged in the last decade as one such promising multidimensional measure of MS disease activity ^13^. On the imaging side, recent technological advances for improved acquisition and analyses are likely to overcome some of the current challenges related to brain scan reproducibility and across-site harmonization and contribute towards a predictive MRI marker for disability and the effect of treatment in patients with MS in a clinical setting ^4^.

Essentially, brain age estimation uses machine learning on a large training set of MRI data from healthy controls (HC) to develop a model that can accurately predict the individual age from brain imaging data ^14–16^. Utilizing sensitive measures of MRI-based brain morphometry, brain age estimation provides a robust imaging-based biomarker with potential to yield novel insights into similarities and differences of disease pathophysiology across brain disorders ^15 17^ Such imaging-based brain age has been shown to be reliable both within and between MRI scanners, and is a candidate biomarker of an individual’s brain health and integrity ^14 15 17^ Different approaches to brain age estimation utilize information from a variety of brain regions (e.g. hippocampus, subcortical, grey matter and white matter ^18^) or MRI sequences (e.g. T1, T2, diffusion tensor imaging and functional MRI) to inform the estimation model ^17^ An older appearing brain, which is related to advanced physiological and cognitive ageing and mortality ^17^, has been found to be an imaging-based hallmark across several brain disorders. To our knowledge, only one preprint (n = 254 MS patients) and one abstract (n = 17 MS patients) have reported brain age estimations in MS ^15 19^, and both reported older appearing brains in patients with MS compared to HC.

Here, combining cross-sectional and sensitive measures of MRI-based regional and global brain morphometry in MS and HC (cross-sectional only), we tested the hypothesis that MS patients have higher estimates of brain age than HC in a cross-sectional design. Next, using longitudinal MRI data in MS patients we tested the hypothesis that brain aging accelerates in MS and that the rate of acceleration is associated with a more severe clinical outcome, investigating the associations between the rate of brain aging, disease-modifying treatment (DMT) and clinical trajectories.

## MATERIALS AND METHODS

### Participants

We included 76 MS patients recruited at Oslo University Hospital ^20 21^. All patients were diagnosed with MS between January 2009 and December 2012 according to the revised McDonald Criteria ^22^ and were enrolled in the study on average 14 months (±11.8) after the date of diagnosis (time point 1). Most patients also participated in two follow-up examinations on average 26 months (±11.7, time point 2, n = 60) and 66 months (±13.3, time point 3, n = 62) after the date of diagnosis. At each visit, all patients completed a neurological examination by a Neurostatus certified medical doctor (http://www.neurostatus.com) within the same week as their MRI scan. DMTs were categorized into the following groups; 0: no treatment; 1: glatiramer acetate, interferons, teriflunomide or dimetylfumarate; and 2: fingolimod, natalizumab or alemtuzumab.

The HC group was recruited through newspaper ads or after a stratified random selection drawn from the Norwegian National Population Registry to two parallel studies ^18 23^. Exclusion criteria included estimated intelligence quotient (IQ) <70, history of neurologic or psychiatric disease and current medication significantly affecting the nervous system ^24^.

The project was approved by the local ethics committee and in line with the Declaration of Helsinki. All participants received oral and written information and gave their written informed consent.

### MRI acquisition

All MS patients were scanned at up to three time points between January 2012 and August 2017, using the same 1.5 T scanner (Avanto, Siemens Medical Solutions; Erlangen, Germany) equipped with a 12-channel head coil. Structural MRI data were collected using a 3D T1-weighted MPRAGE (Magnetization Prepared Rapid Gradient Echo) sequence, with the following parameters: repetition time (TR) / echo time (TE) / flip angle / voxel size / field of view (FOV) / slices / scan time / matrix / time to inversion = 2400 ms / 3.61 ms / 8° / 1.20 × 1.25 × 1.25 mm / 240 / 160 sagittal slices / 7:42 minutes / 192 × 192 / 1000 ms. The MRI sequence was kept identical during the scanning period. FLAIR (Fluid attenuation inversion recovery), T2 and pre-and post-gadolinium 3D T1 sequences were attained and used for neuroradiological evaluation ^20^.

Fifty-eight of the MS patients were also scanned at Oslo University Hospital on a 3 T GE 750 Discovery MRI scanner with a 32-channel head coil at time point 3 between August 2016 and June 2017 during the same week they were scanned at the 1.5 T scanner for time point 3. HCs were scanned solely on the 3 T scanner. Structural MRI data were collected using a 3D high-resolution IR-prepared fast spoiled gradient echo (FSPGR) T1-weighted sequence (3D BRAVO) with the following parameters: TR / TE / flip angle / voxel size / FOV / slices / scan time = 8.16 ms / 3.18 ms / 12° / 1 × 1 × 1 mm / 256 × 256 mm / 188 sagittal slices / 4:42 minutes.

### MRI pre- and postprocessing

Using the T1-weighted scans we performed cortical reconstruction and volumetric segmentation with FreeSurfer 5.3 (http://surfer.nmr.mgh.harvard.edu/) ^25^. To extract reliable volume and thickness estimates, images included in the longitudinal 1.5 T MRI dataset were processed with the longitudinal stream in FreeSurfer ^26^. Specifically an unbiased within-subject template space and image was created using robust, inverse consistent registration ^27^. Several processing steps, such as skull stripping, Talairach transforms, atlas registration as well as spherical surface maps and parcellations were then initialized with common information from the within-subject template, increasing reliability and power ^26^.

Manual quality control of the MRI scans from patients was performed by trained research personnel to identify and edit segmentation errors where possible (n = 43 MRI scans) and exclude data of insufficient quality (n = 6 MRI scans). In addition, eight brain scans were removed due to missing sequences of the 263 MRI scans from MS patients. Lesion filling was performed utilizing automatically generated lesion masks from Cascade ^28^ with the lesion filling tool (https://fsl.fmrib.ox.ac.uk/fsl/fslwiki/lesion_filling) in FSL ^29^. The lesion masks were assessed by a trained neuroradiologist. For a probabilistic representation of the lesions, the lesion masks were normalized into MNI-standard space using FSL FLIRT ^30^, with the corresponding T1 image as an intermediate. A probabilistic representation of the lesions across all patients is shown in Supplementary Fig. 1.

### Brain age estimation model

The training set for the brain age estimation included data from 3208 HC >12 years (54 % women, mean age 47.5 (±19.8), age range 12-95) obtained from several publicly available datasets (Supplementary Fig. 2). We trained one “xgboost” machine learning model for each sex to predict brain age derived from a total of 1118 cortical and subcortical brain imaging features ^15 31 32^.

Based on a recent implementation ^15^, brain age estimations were performed both using global and regional features as input. The brain regions reported were based on lobe labels from Freesurfer ^25^. The brain age estimation model was carefully evaluated using 10-fold cross-validation within the training set and showed good performance and generalizability (Supplementary Fig. 3, r = 0.91). For all patients and HC, we calculated the brain age gap (BAG, defined as the difference between chronological age and imaging-based brain age). Using linear regressions, we removed any common variance with age, age^2^ and sex to account for confounding factors before submitting the residualized version of BAG to further analyses ^33^. When pooling estimates of BAG from the 1.5 T and 3 T scanners, we adjusted BAG for scanner effect on BAG estimates by extracting the scanner coefficient from a linear mixed effects (LME) model for global and all brain regions. When comparing BAG between patients and matched HCs we report the actual adjusted difference in BAG between these two groups.

### Statistical analyses

We used R (R Core Team, Vienna, 2018) for statistical analyses. All LME models accounted for age, age^2^, sex and scanner ^34^.

We estimated an annualized rate of change in BAG by dividing the total change in BAG by the time interval between the time points. We utilized the longest time interval between time points and excluded MS patients lacking longitudinal data (n = 8). The unit used for brain aging is “change in BAG per year”. Here, 0 would indicate that the rate of brain aging corresponds to chronological aging, and positive and negative values correspond to accelerated and decelerated brain aging compared to chronological aging, respectively. For all brain regions we tested for significant change in the rate of brain aging by performing a one-sample t-test on BAG with 0 set as test value. To assess brain aging within groups stratified by DMT, we used the coefficients from linear models with the individual slope in BAG as dependent variable and age, age^2^, sex and DMT group as independent variables. We estimated the annual global brain atrophy by comparing estimated total brain volume from the Freesurfer output (BrainSegVolNotVent) between time points.

To assess reliability of brain age across time we computed the intraclass correlation coefficient (ICC) using the R package “irr” (https://CRAN.R-project.org/package=irr). Figures were made using “ggplot2” ^35^ and “cowplot” (https://CRAN.R-project.org/package=cowplot) in R. To control for multiple testing we adjusted the p-values using false discovery rate (FDR) ^36^ procedures implemented in the R package “p.adjust” (http://stat.ethz.ch/R-manual/R-devel/library/stats/html/p.adjust.html). The linear mixed effect models were performed using the R package “nlme” (https://CRAN.R-project.org/package=nlme).

### Data availability

The data are not publicly available due to local restrictions, since they contain information that may compromise the privacy of research participants. The code needed to reproduce our results is available from the authors upon request.

## RESULTS

### Participant demographics and characteristics

Table 1 summarizes the demographic and clinical characteristics of all MS patients. Key demographic variables regarding HC are summarized in Supplementary Table 1. The majority of the MS patients were women (71%) and mean age at inclusion was 34.8 years (±7.2). On average they were examined 1.2, 2.2 and 5.5 years after diagnosis. Most patients used first line treatment; 65%, 48% and 37% at time point 1, 2 and 3, respectively. Second line treatments were used by 13%, 23% and 32% of the MS patients at time point 1, 2 and 3, respectively. At time point 2 and 3, 53% and 44% of the patients were categorized as NEDA-3 (no clinical progression, no new lesions observable in MRI and no new attacks).

**Table 1.**
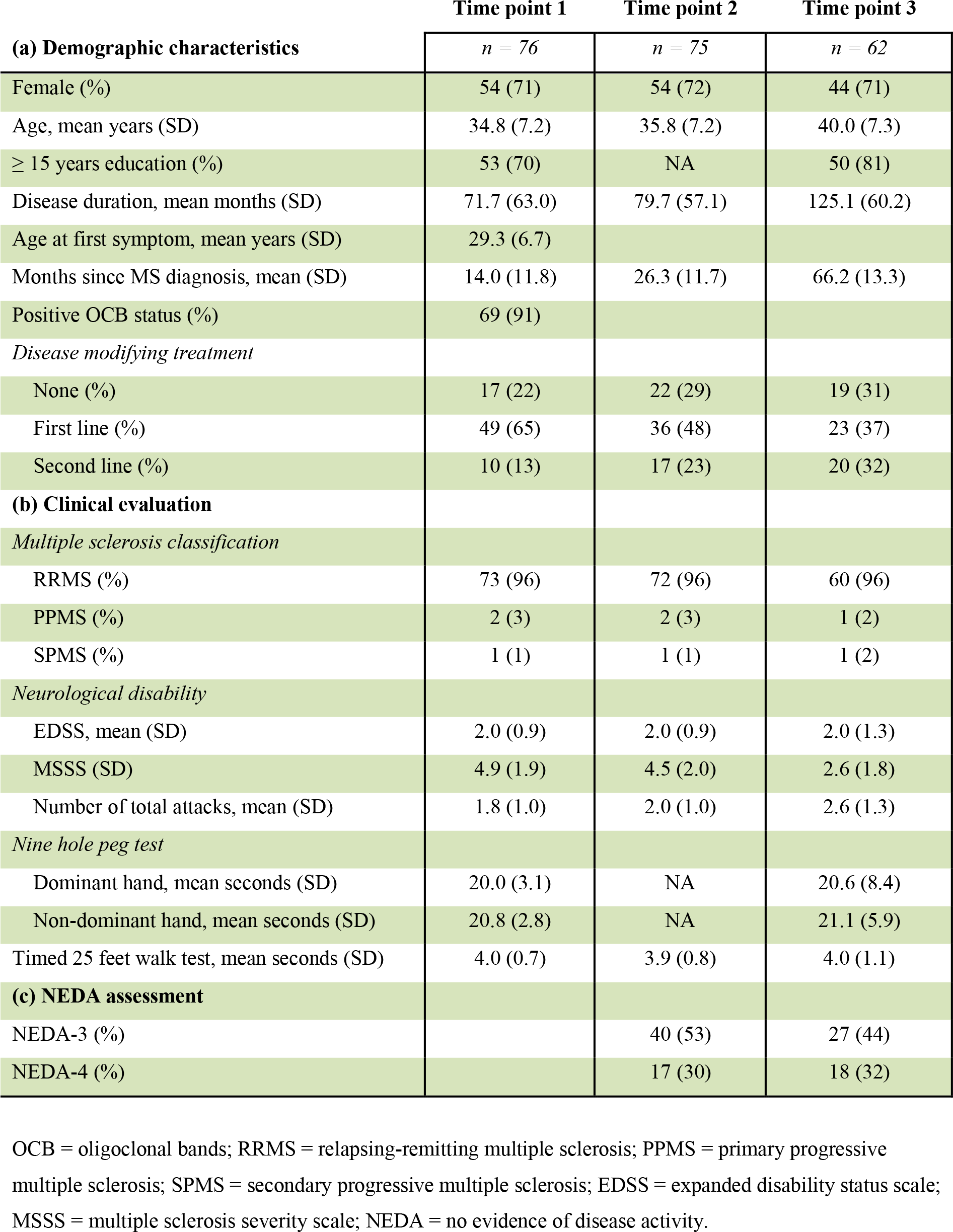
Demographic and clinical characteristics of the multiple sclerosis patients.

### Cross-sectional case-control analyses (3 T)

At time point 3 we found significantly higher BAG for the MS cohort compared to matched HC for all brain regions except the temporal region (Fig. 1; Table 2). The most prominent differences in BAG were 4.4 years for global BAG (Cohen’s D = 0.69) and 6.2 years for subcortical and cerebellar brain regions (Cohen’s D = 0.72).

At time point 3, 58 MS patients underwent one MRI scanning in the 1.5 T and one in the 3 T scanner with two days apart. Whereas absolute estimates of brain age varied between scanners for all brain regions except insula (BAG scanner difference −6.08 to 10.60 years, see Supplementary Table 2; Supplementary Fig. 4), brain age estimates from the two scanners were highly correlated for global BAG and all brain regions (r = 0.67-0.86, p<0.001).

**Figure 1.**
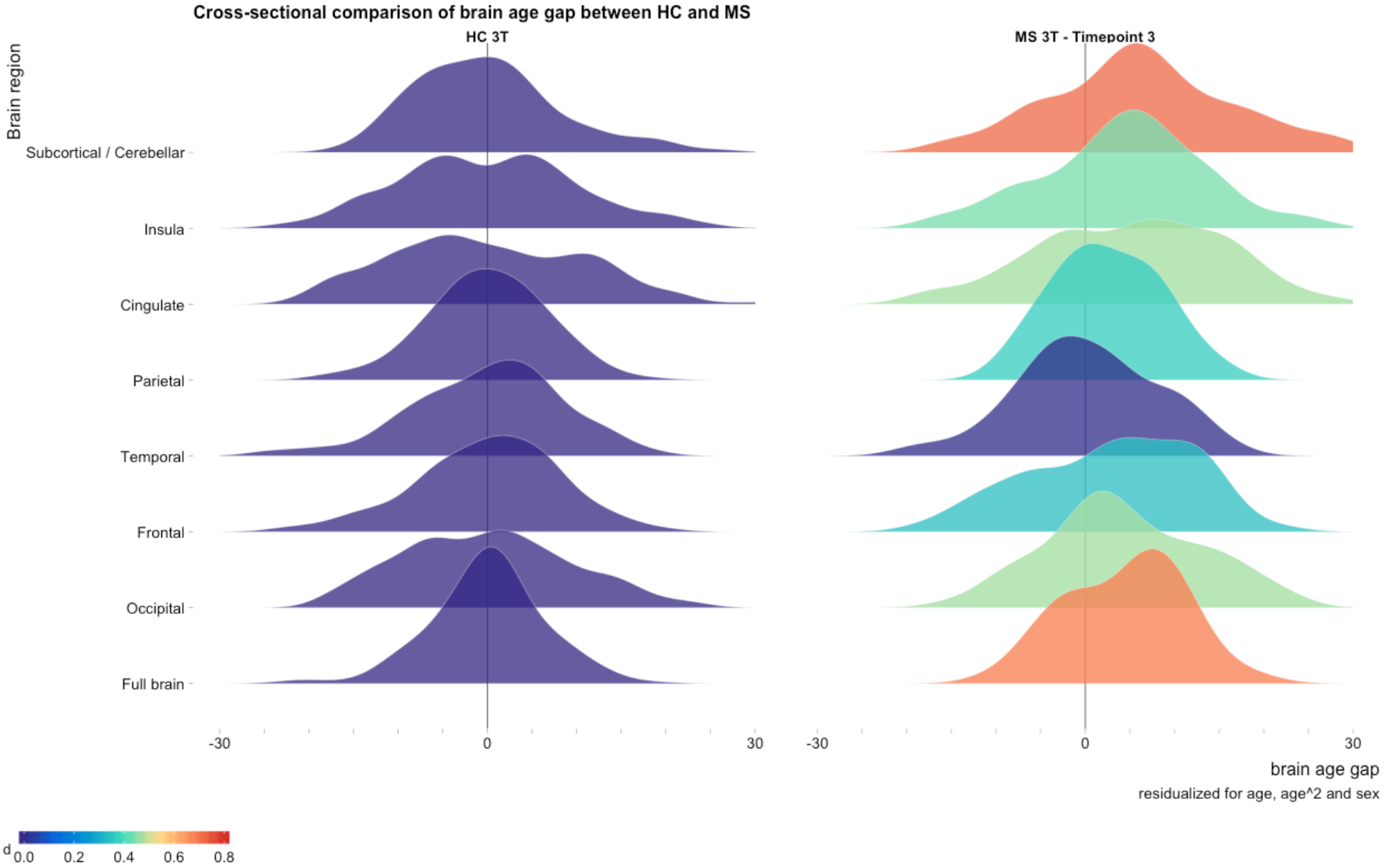
Cross-sectional comparison of brain age gap between MS patients and healthy controls. The distribution of brain age gaps across brain regions based on the cross-sectional 3 T MRI data from matched HC and MS patients at time point 3. We found increased brain age gaps for all brain regions except from the temporal brain region. Brain age gaps are residualized for age, age^2^ and sex. Cohen’s D effect sizes for the brain age gap between HC and MS patients are depicted using the colour bar.

### Longitudinal MS sample (1.5 T)

Fig. 2 shows the estimated brain age and rate of brain aging within the longitudinal MS cohort. The correlations between chronological age and global brain age were r = 0.71 for time point 1, r = 0.70 for time point 2 and r = 0.69 for time point 3. After adjusting for scanner effects mean global BAG was 2.8 (±9.0) for time point 1, 3.3 (±9.4) for time point 2 and 4.6 (±9.8) for time point 3 in the longitudinal MS sample (Fig. 2B, Table 2 and Supplementary table 3).

We found a significant annual increase in global BAG of 0.41 (±1.23) years (p = 0.008) in patients with MS (Fig. 3; Supplementary Table 4 and 5). None of the regional measures showed significantly increased annual change in BAG (Supplementary Table 5).

We found no significant difference in BAG between the raw and the lesion filled MRI scans, and the two versions yielded high correlation in BAG (r = 0.98). Data processed with the longitudinal stream in FreeSurfer had significantly lower BAG than the cross-sectionally processed MRI scans (difference in BAG 4.9 years, p<0.001) and lesion filled MRI scans (difference in BAG 5.1 years, p = <0.001) (Supplementary Fig. 5 and 6; Supplementary Table 6 and 7).

ICCs for all brain regions across all time points varied from 0.79-0.94 for residualized brain age gap and 0.78-0.95 for raw predicted age. The regional feature of cerebellar and subcortical brain regions showed highest reliability with an ICC of 0.94 for BAG and 0.95 for predicted age (Supplementary Table 8).

Mean annualized estimated change in global brain volume from all three time points. from Freesurfer was −0.30 % (±0.53 %). ICC for global brain volume was 0.97-0.99. Mean annualized change in WMLL was 504 mm^3^ (±1299 mm^3^). ICC for WMLL was 0.93-0.99.

**Figure 2.**
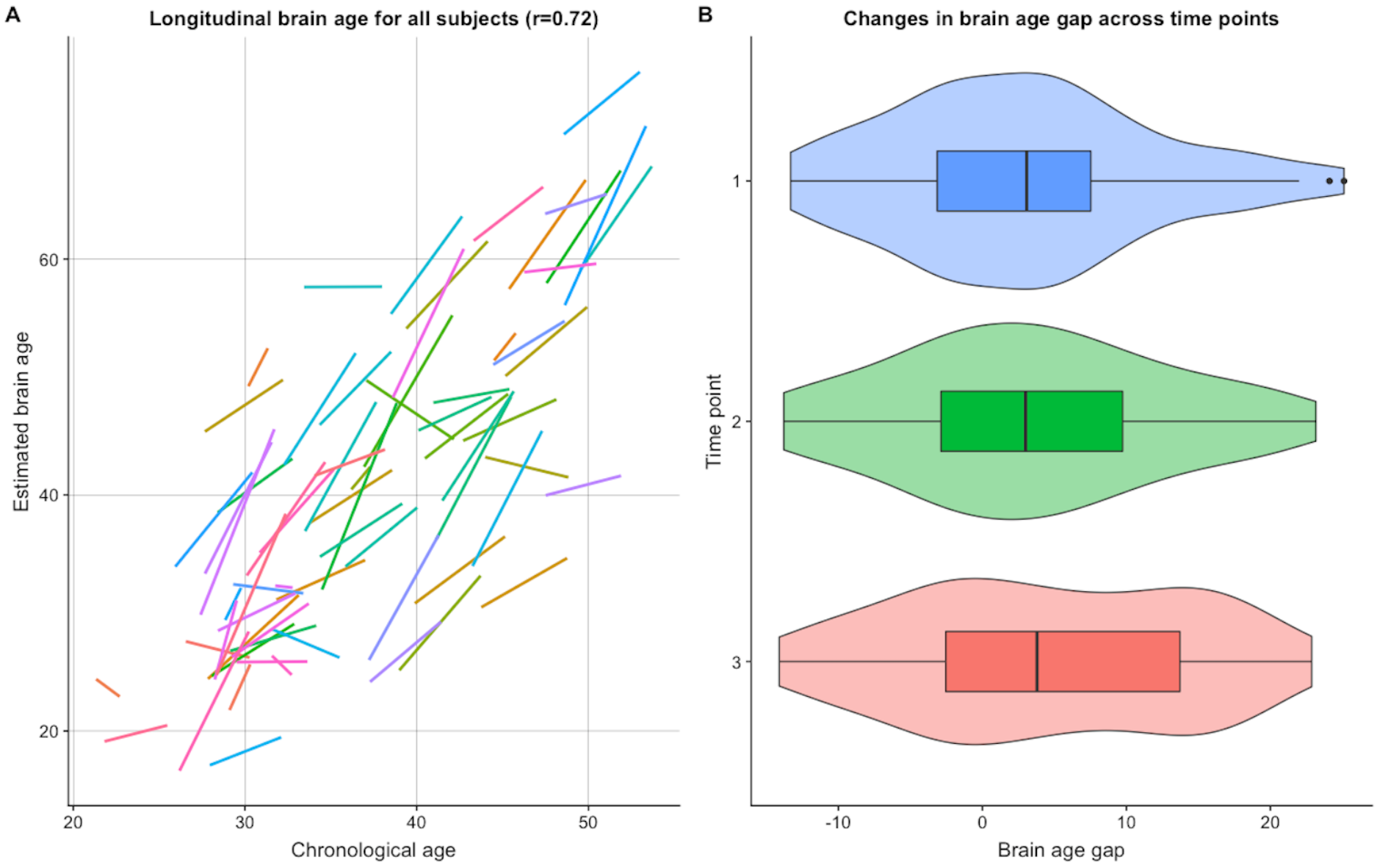
Visualization of brain age in the longitudinal MS cohort. **A,** MS subjects are depicted with linear brain age slopes using linear regression models to visualize individualised estimations of brain aging. Only participants with more than one MRI scan are included (n = 68). Mean annual increase in global BAG was 0.41(±1.23) years (p = 0.008) in patients with MS. **B**, Difference between chronological and predicted age (brain age gap) are shown for all three time points separately. After adjusting for scanner effects mean brain age gap was 2.8 (±9.0) for time point 1, 3.3 (±9.4) for time point 2 and 4.6 (±9.8) for time point 3 in the longitudinal sample. The distributions of the brain age estimates are visualized using box and violin plots.

**Figure 3.**
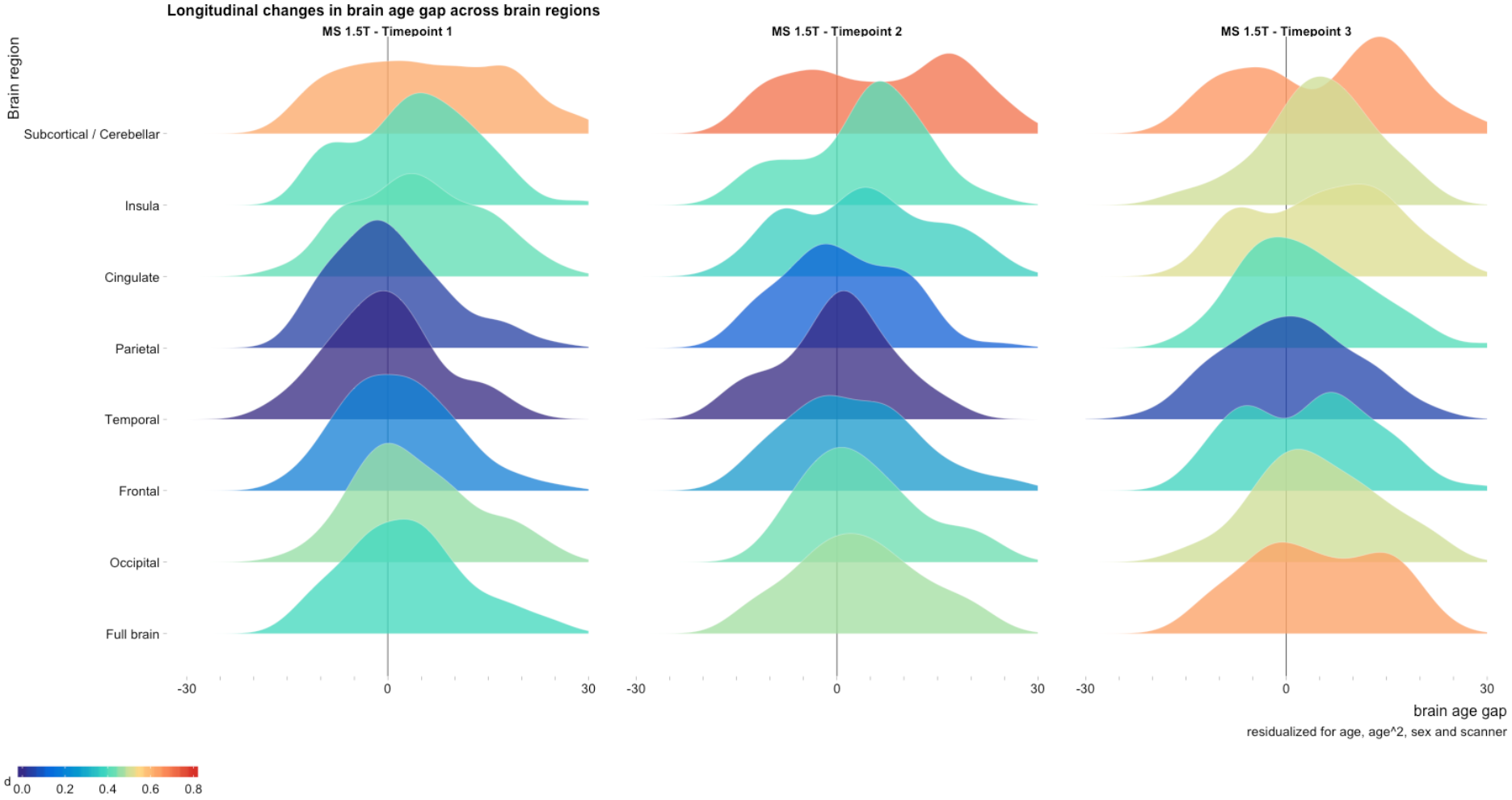
Longitudinal changes in brain age gap across brain regions. The distribution of brain age gaps across brain regions based on the longitudinal 1.5 T MRI sample. Brain age gaps from the MS sample are compared with the cross-sectional 3 T HC sample and residualized for age, age^2^, sex and scanner. The full brain estimates showed a significant accelerated rate of brain aging compared to chronological aging (annual increase in brain age gap 0.41 (p = 0.008)). Cohen’s D effect sizes for the brain age gap between MS and HC are depicted using the colour bar.

**Table 2.**
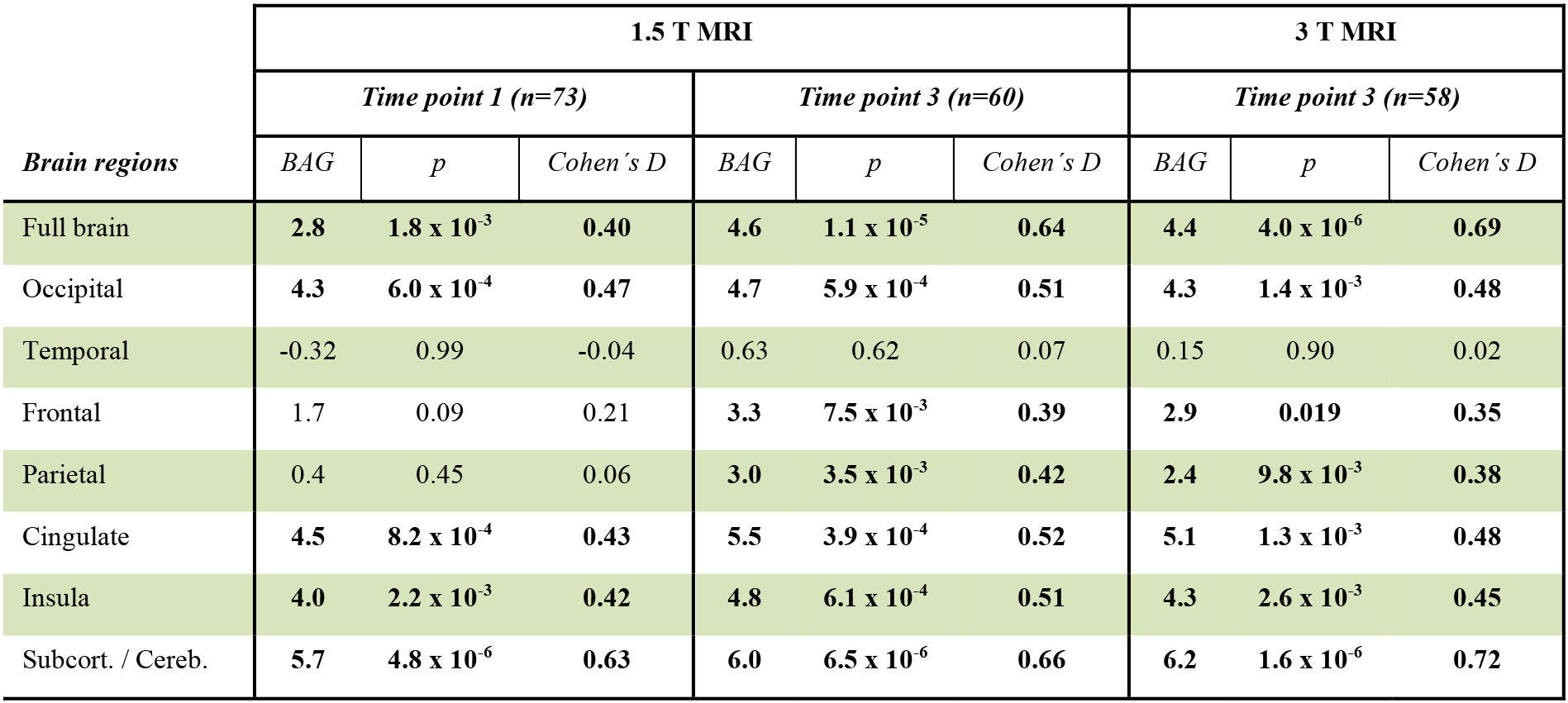
Differences in brain age gap between the multiple sclerosis patients and healthy controls (n = 235).

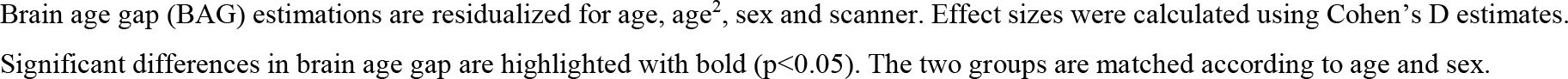

### Associations between global brain age and clinical outcomes

Table 3 (BAG) and Table 4 (annual rate of brain aging) show summary statistics from the multiple regressions testing for associations with demographic, clinical and MRI variables in the longitudinal MS cohort. After accounting for multiple testing, significant associations were found between BAG and brain atrophy (Cohen’s D = −0.07, p = 0.01) and WMLL (Cohen’s D = −1.23, p = 3.0 × 10^−4^), indicating higher BAG at baseline with higher WMLL and increased brain atrophy. For longitudinal estimates of brain aging we found significant associations with brain atrophy (Cohen’s D = 0.86, p = 4.3 × 10^−15^) and change in WMLL (Cohen’s D = 0.55, p = 0.015), indicating higher rates of brain aging in patients with higher levels of brain atrophy and more progressive changes in WMLL. WMLL also showed a significant correlation with BAG for cerebellar and subcortical regions (Cohen’s D = −1.23, p = 3.2 × 10^−3^).

In the longitudinal data, One Sample t-test revealed a significant increase in BAG in DMT group 0 (0.92 (±0.82), p = 5.4×10^−4^), and no significant changes in DMT groups 1 (0.13 (±1.3), p = 0.63) and 2 (0.35 (±1.3), p = 0.26). However, linear models and a LME analysis revealed no significant group differences in the rate of brain aging (f-value = 2.47, p = 0.09) (Supplementary Fig. 7).

**Table 3.**
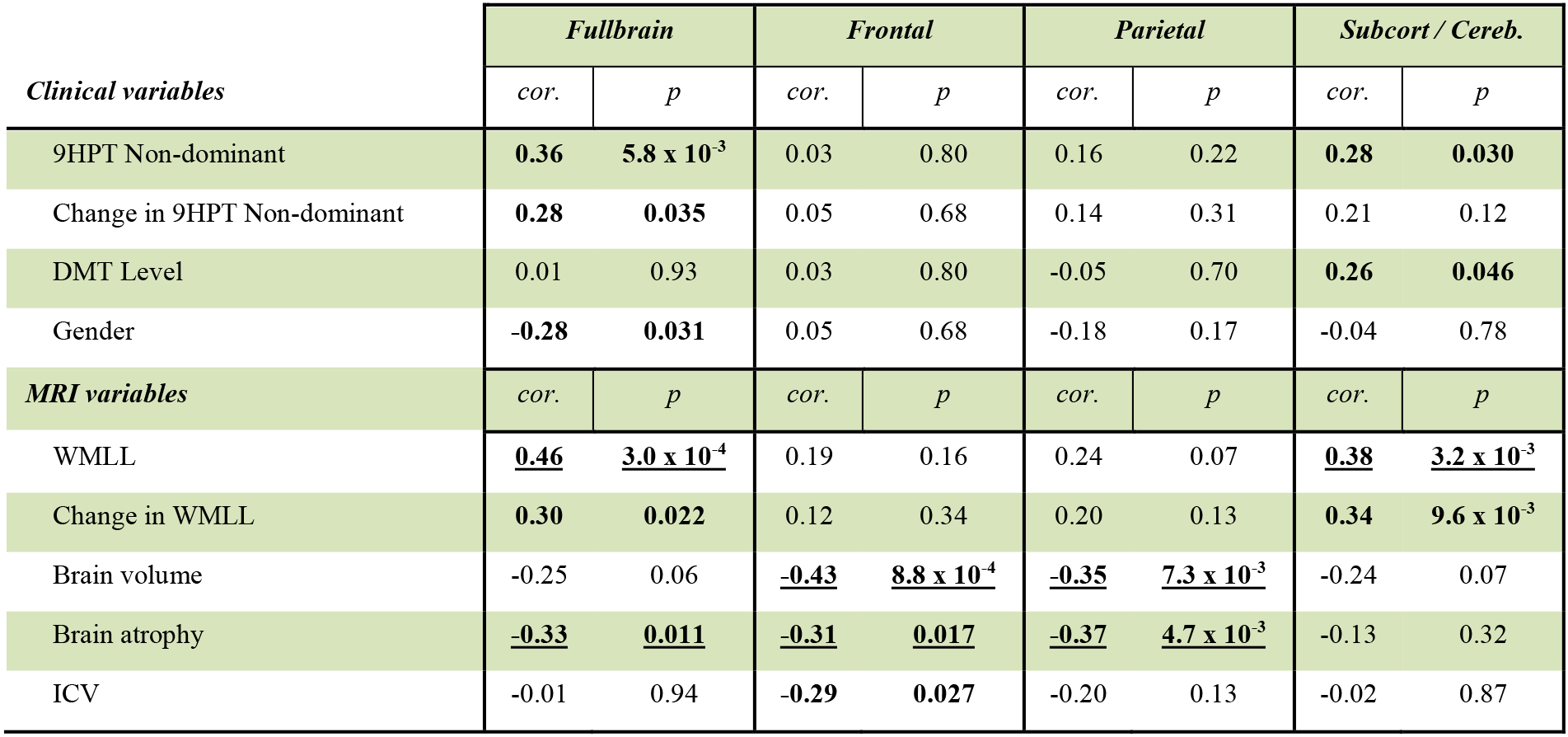
Pearson’s correlations between brain age gap and relevant clinical and MRI variables.

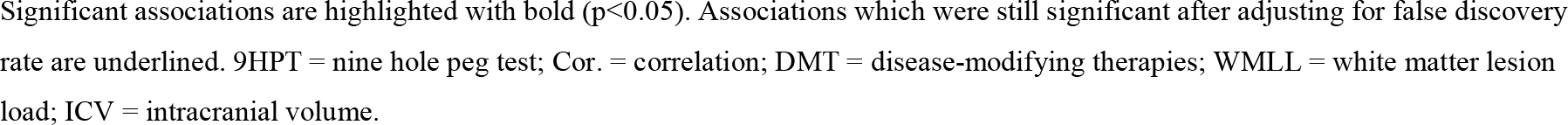

**Table 4.**
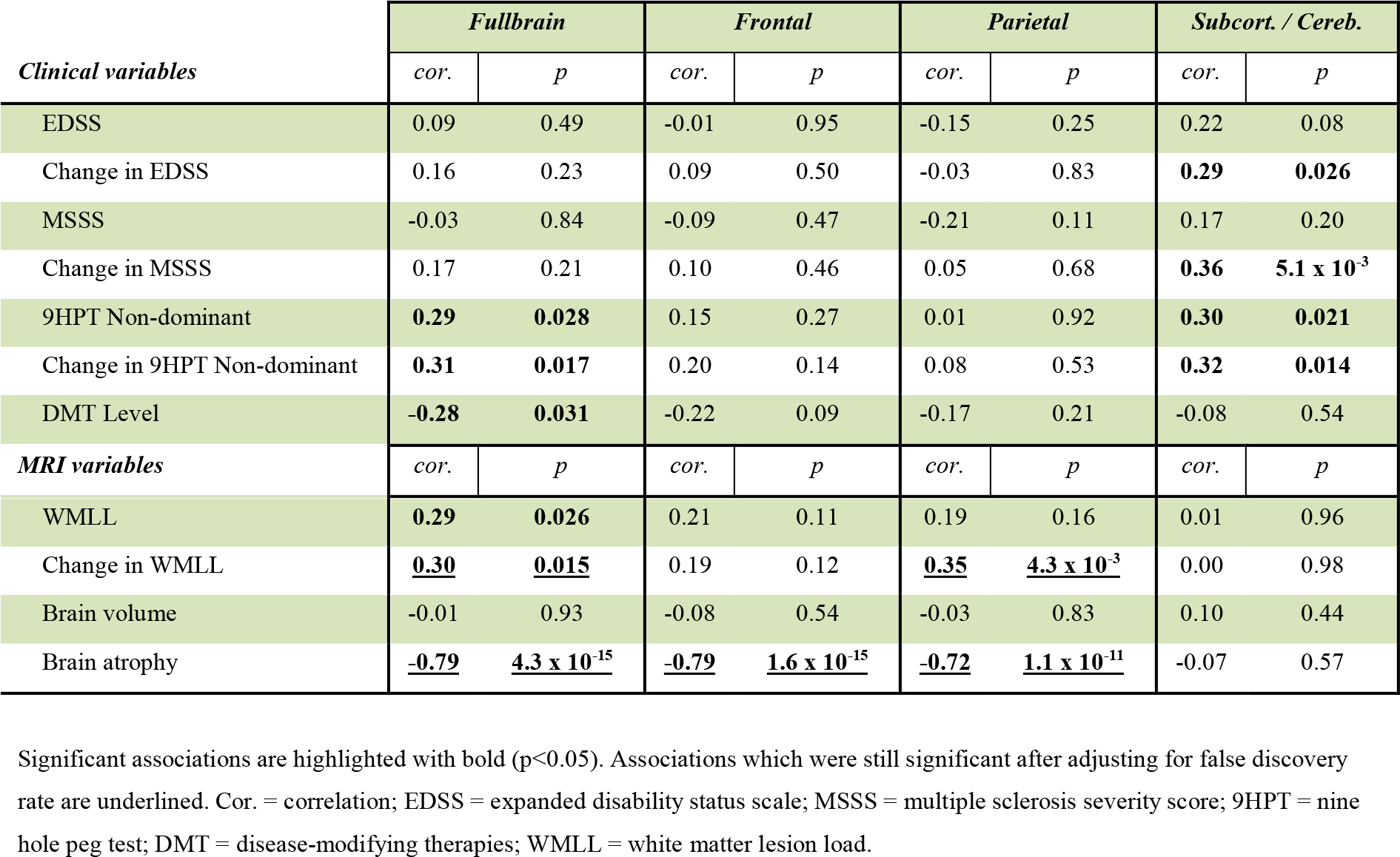
Pearson’s correlations between annual rate of brain aging and relevant clinical and MRI variables on time point 3.

| *Clinical variables* | *Fullbrain* |  | *Frontal* |  | *Parietal* |  | *Subcort. / Cereb.* |  |
| --- | --- | --- | --- | --- | --- | --- | --- | --- |
|  | *cor.* | *p* | *cor.* | *p* | *cor.* | *p* | *cor.* | *p* |
| EDSS | 0.09 | 0.49 | -0.01 | 0.95 | -0.15 | 0.25 | 0.22 | 0.08 |
| Change in EDSS | 0.16 | 0.23 | 0.09 | 0.50 | -0.03 | 0.83 | **0.29** | **0.026** |
| MSSS | -0.03 | 0.84 | -0.09 | 0.47 | -0.21 | 0.11 | 0.17 | 0.20 |
| Change in MSSS | 0.17 | 0.21 | 0.10 | 0.46 | 0.05 | 0.68 | **0.36** | **$5.1 \times 10^{-3}$** |
| 9HPT Non-dominant | **0.29** | **0.028** | 0.15 | 0.27 | 0.01 | 0.92 | **0.30** | **0.021** |
| Change in 9HPT Non-dominant | **0.31** | **0.017** | 0.20 | 0.14 | 0.08 | 0.53 | **0.32** | **0.014** |
| DMT Level | **-0.28** | **0.031** | -0.22 | 0.09 | -0.17 | 0.21 | -0.08 | 0.54 |
| *MRI variables* | *cor.* | *p* | *cor.* | *p* | *cor.* | *p* | *cor.* | *p* |
| WMLL | **0.29** | **0.026** | 0.21 | 0.11 | 0.19 | 0.16 | 0.01 | 0.96 |
| Change in WMLL | <u>**0.30**</u> | <u>**0.015**</u> | 0.19 | 0.12 | <u>**0.35**</u> | <u>**$4.3 \times 10^{-3}$**</u> | 0.00 | 0.98 |
| Brain volume | -0.01 | 0.93 | -0.08 | 0.54 | -0.03 | 0.83 | 0.10 | 0.44 |
| Brain atrophy | <u>**-0.79**</u> | <u>**$4.3 \times 10^{-15}$**</u> | <u>**-0.79**</u> | <u>**$1.6 \times 10^{-15}$**</u> | <u>**-0.72**</u> | <u>**$1.1 \times 10^{-11}$**</u> | -0.07 | 0.57 |

Significant associations are highlighted with bold ( $p < 0.05$ ). Associations which were still significant after adjusting for false discovery rate are underlined. Cor. = correlation; EDSS = expanded disability status scale; MSSS = multiple sclerosis severity score; 9HPT = nine hole peg test; DMT = disease-modifying therapies; WMLL = white matter lesion load.

## DISCUSSION

Here, by using advanced cross-sectional and longitudinal MRI-based neuroimaging data as basis for brain age estimation based on machine learning, we tested the hypotheses that patients with MS on average show higher brain age than HC, and that the rate of brain aging is affected by treatment. We found an accelerated brain aging in MS patients compared to controls by cross-sectional and longitudinal brain scans and suggested that increased rates of longitudinal brain aging are associated with higher rates of brain atrophy and increasing WMLL.

We report that MS patients on average had 4.4 years higher BAG compared to HC (Cohen’s D = 0.68). The present findings are in line with preliminary findings from Kaufmann and colleagues ^15^. Other studies on this topic are warranted. To our knowledge, results of other comparable studies are not yet available.

For subcortical and cerebellar brain regions we found a higher BAG in MS compared with HC (BAG 6.2 years, Cohen’s D = 0.72), which was already evident at time point 1 (BAG 5.7 years, Cohen’s D = 0.63). The regional variation observed in the BAG estimates in our study may reflect different affinity of MS pathology in different brain regions and aligns with previously shown lesion probability maps in MS ^4 37^.

In our longitudinal patient sample, the annual rate of brain aging for global BAG exceeded that of chronological aging by 0.41 years per year (p = 0.008), which may partly be explained by chronic inflammatory processes that drive neurodegeneration in MS ^3^. The association between BAG and brain aging, brain volume and brain atrophy were expected since our estimation model was based on regional and global structural MRI features.

Of notice, brain age estimation takes into account subtle and regional changes that are not included in the global brain atrophy measure. The associations between BAG and brain aging, brain atrophy, brain volume, WMLL and change in WMLL showed significant and regional differences. The associations between change in WMLLs and brain aging were significant for occipital, temporal and parietal brain regions in addition to global BAG (Supplementary Table 9). This shows that regional brain age estimation can capture regional specificity of MS pathology, which is in line with the observations that MS lesions show regional affinity ^14 15 17 37^

The significant increase in BAG in the patients receiving no treatment warrants further investigations but is of clinical interest. However, the lack of randomization and group differences in the subgroups calls for caution in the interpretations of this relative small dataset.

The preliminary findings reported by Kaufmann et al. revealed a significant association between BAG and EDSS (Fisher z = 0.23) ^15^, i.e. that patients with older appearing brains have a higher clinical disease burden. In the current study, multiple regression analyses revealed nominally significant (p<0.05, uncorrected) associations between some clinical, cognitive and imaging variables and BAG as well as brain aging for specific brain regions (Supplementary Table 9 and 10). However, these associations did not survive correction for multiple testing, and further studies are needed to assess the robustness of these observations.

Some limitations should be considered when interpreting the results. First, although the cross-sectional case-control comparison and the within-patient longitudinal analysis jointly suggest accelerated brain aging in patients with MS, a longitudinal sample of HCs would have enabled us to directly compare the rate of brain aging between patients and controls. Next, the current brain age model was exclusively based on gross morphometric features. The MS patients were naturalistic, hence any results based on DMT status must be interpreted with care.

In conclusion, using advanced cross-sectional imaging data and machine learning methods we report that patients with MS show evidence of increased brain aging compared to healthy controls. In the longitudinal data we found that MS patients have accelerated brain aging. Higher rates of longitudinally measured brain aging were associated with higher levels of brain atrophy and longitudinal progression of changes in WMLL. These results show that brain age estimation is a promising and intuitive tool for monitoring of the individual disease course in MS and may guide a personalized treatment approach.

## ACKNOWLEDGMENTS

We thank all the patients who participated in our study. We acknowledge the collaboration with members of the Multiple Sclerosis Research Group at the University of Oslo and Oslo University hospital, especially Professor Elisabeth G. Celius.

## SUPPLEMENTARY MATERIAL

Supplementary material is included in separate files.

## REFERENCES

1. Berg-Hansen P, Moen SM, Harbo HF, et al. High prevalence and no latitude gradient of multiple sclerosis in Norway. Mult Scler 2014;20(13):1780–2. doi:10.1177/1352458514525871 [published Online First: 2014/03/08]

2. Noseworthy JH, Lucchinetti C, Rodriguez M, et al. Multiple Sclerosis. The New England Journal of Medicine 2000;343(13):938–52. doi:10.1056/NEJM200009283431307

3. Friese MA, Schattling B, Fugger L. Mechanisms of neurodegeneration and axonal dysfunction in multiple sclerosis. Nat Rev Neurol 2014;10(4):225–38. doi:10.1038/nrneurol.2014.37 [published Online First: 2014/03/19]

4. Rocca MA, Comi G, Filippi M. The Role of T1-Weighted Derived Measures of Neurodegeneration for Assessing Disability Progression in Multiple Sclerosis. Front Neurol 2017;8:433. doi:10.3389/fneur.2017.00433

5. Filippi M, Preziosa P, Rocca MA. MRI in multiple sclerosis: what is changing? Curr Opin Neurol 2018;31(4):386–95. doi:10.1097/WG3.0000000000000572 [published Online First: 2018/06/29]

6. Filippi M, Preziosa P, Rocca MA. Brain mapping in multiple sclerosis: Lessons learned about the human brain. Neuroimage 2017 doi:10.1016/j.neuroimage.2017.09.021

7. Filippi M, Rocca MA, Ciccarelli O, et al. MRI criteria for the diagnosis of multiple sclerosis: MAGNIMS consensus guidelines. Lancet Neurol 2016;15(3):292–303. doi:10.1016/s1474-4422(15)00393-2 [published Online First: 2016/01/30]

8. Altmann DR, Jasperse B, Barkhof F, et al. Sample sizes for brain atrophy outcomes in trials for secondary progressive multiple sclerosis. Neurology 2009;72(7):595–601. doi:10.1212/01.wnl.0000335765.55346.fc

9. Popescu V, Klaver R, Versteeg A, et al. Postmortem validation of MRI cortical volume measurements in MS. Hum Brain Mapp 2016;37(6):2223–33. doi:10.1002/hbm.23168

10. Chard D, Trip SA. Resolving the clinico-radiological paradox in multiple sclerosis. F1000Res 2017;6:1828. doi:10.12688/f1000research.11932.1

11. Kurtzke FJ. Rating neurologic impairment in multiple sclerosis: An expanded disability status scale (EDSS). Neurology 1983;33(11):1444–52. doi:10.1212/WNL.33.11.1444

12. van Munster CE, Uitdehaag BM. Outcome Measures in Clinical Trials for Multiple Sclerosis. CNS Drugs 2017;31(3):217–36. doi:10.1007/s40263-017-0412-5 [published Online First: 2017/02/12]

13. Giovannoni G, Bermel R, Phillips T, et al. A brief history of NEDA. Mult Scler Relat Disord 2018;20:228–30. doi:10.1016/j.msard.2017.07.011 [published Online First: 2018/03/27]

14. Cole JH, Poudel RPK, Tsagkrasoulis D, et al. Predicting brain age with deep learning from raw imaging data results in a reliable and heritable biomarker. Neuroimage 2017 doi:10.1016/j.neuroimage.2017.07.059

15. Kaufmann T, van der Meer D, Doan NT, et al. Genetics of brain age suggest an overlap with common brain disorders. bioRxiv 2018 doi:10.1101/303164

16. Franke K, Ziegler G, Klöppel S, et al. Estimating the age of healthy subjects from T1-weighted MRI scans using kernel methods: Exploring the influence of various parameters. Neuroimage 2010;50(3):883–92. doi:https://doi.org/10.1016Zj.neuroimage.2010.01.005

17. Cole JH, Franke K. Predicting Age Using Neuroimaging: Innovative Brain Ageing Biomarkers. Trends Neurosci 2017;40(12):681–90. doi:10.1016/j.tins.2017.10.001

18. Richard G, Kolskaar K, Sanders A-M, et al. Assessing distinct patterns of cognitive aging using tissue-specific brain age prediction based on diffusion tensor imaging and brain morphometry. bioRxiv 2018 doi:10.1101/313015

19. Raffel J, Cole J, Record C, et al. Brain Age: A novel approach to quantify the impact of multiple sclerosis on the brain (P1.371). Neurology 2017;88(16 Supplement)

20. Nygaard GO, Walhovd KB, Sowa P, et al. Cortical thickness and surface area relate to specific symptoms in early relapsing-remitting multiple sclerosis. Mult Scler 2015;21(4):402–14. doi:10.1177/1352458514543811 [published Online First: 2014/08/21]

21. Nygaard GO, Celius EG, de Rodez Benavent SA, et al. A Longitudinal Study of Disability, Cognition and Gray Matter Atrophy in Early Multiple Sclerosis Patients According to Evidence of Disease Activity. PLoS One 2015;10(8):e0135974. doi:10.1371/journal.pone.0135974 [published Online First: 2015/08/19]

22. Polman CH, Reingold SC, Banwell B, et al. Diagnostic criteria for multiple sclerosis: 2010 revisions to the McDonald criteria. Ann Neurol 2011;69(2):292–302. doi:10.1002/ana.22366 [published Online First: 2011/03/10]

23. Moberget T, Doan NT, Alnaes D, et al. Cerebellar volume and cerebellocerebral structural covariance in schizophrenia: a multisite mega-analysis of 983 patients and 1349 healthy controls. Mol Psychiatry 2018;23(6):1512–20. doi:10.1038/mp.2017.106 [published Online First: 2017/05/17]

24. Doan NT, Engvig A, Zaske K, et al. Distinguishing early and late brain aging from the Alzheimer’s disease spectrum: consistent morphological patterns across independent samples. Neuroimage 2017;158:282–95. doi:10.1016/j.neuroimage.2017.06.070 [published Online First: 2017/07/02]

25. Dale AM, Fischl B, Sereno MI. Cortical surface-based analysis. I. Segmentation and surface reconstruction. Neuroimage 1999;9(2):179–94. doi:10.1006/nimg.1998.0395 [published Online First: 1999/02/05]

26. Reuter M, Schmansky NJ, Rosas HD, et al. Within-subject template estimation for unbiased longitudinal image analysis. Neuroimage 2012;61(4):1402–18. doi:10.1016/j.neuroimage.2012.02.084 [published Online First: 2012/03/21]

27. Reuter M, Rosas HD, Fischl B. Highly accurate inverse consistent registration: a robust approach. Neuroimage 2010;53(4):1181–96. doi:10.1016/j.neuroimage.2010.07.020 [published Online First: 2010/07/20]

28. Damangir S, Manzouri A, Oppedal K, et al. Multispectral MRI segmentation of age related white matter changes using a cascade of support vector machines. J Neurol Sci 2012;322(1-2):211–6. doi:10.1016/j.jns.2012.07.064

29. Jenkinson M, Beckmann CF, Behrens TE, et al. Fsl. NeuroImage 2012;62(2):782–90. doi:10.1016/j.neuroimage.2011.09.015

30. Jenkinson M, Bannister P, Brady M, et al. Improved optimization for the robust and accurate linear registration and motion correction of brain images. Neuroimage 2002;17(2):825–41. [published Online First: 2002/10/16]

31. XGBoost: A Scalable Tree Boosting System. KDD ’16; 2016; San Francisco, California, US. ACM.

32. Glasser MF, Coalson TS, Robinson EC, et al. A multi-modal parcellation of human cerebral cortex. Nature 2016;536(7615):171–78. doi:10.1038/nature18933 [published Online First: 2016/07/21]

33. Le TT, Kuplicki R, McKinney BA, et al. A nonlinear simulation framework supports adjusting for age when analyzing BrainAGE. bioRxiv 2018 doi:10.1101/377648

34. Bernal-Rusiel JL, Greve DN, Reuter M, et al. Statistical analysis of longitudinal neuroimage data with Linear Mixed Effects models. Neuroimage 2013;66:249–60. doi:10.1016/j.neuroimage.2012.10.065

35. Wickham H. ggplot2: Elegant Graphics for Data Analysis: Springer–Verlag New York 2016.

36. Benjamini Y, Hochberg Y. Controlling the False Discovery Rate: A Practical and Powerful Approach to Multiple Testing. Journal of the Royal Statistical Society Series B (Methodological) 1995;57(1):289–300.

37. Gourraud PA, Sdika M, Khankhanian P, et al. A genome-wide association study of brain lesion distribution in multiple sclerosis. Brain 2013;136(Pt 4):1012–24. doi:10.1093/brain/aws363 [published Online First: 2013/02/16]

